# Systematic design of multiplex methylation-specific qPCR for cancer related biomarkers

**DOI:** 10.1101/2024.12.09.627654

**Authors:** Joy Wang, Michelle Yiru Lee, Brian Boyuan Lin, Annie X. Wu

**Author notes:** These authors contributed equally to this work (Joy Wang, Michelle Yiru Lee, Brian Boyuan Lin).

## Abstract

Analysis of circulating cell-free DNA (cfDNA) methylation abnormalities has emerged as a powerful strategy for detecting various diseases, particularly cancers. This study demonstrates a process for developing a multiplex methylation-specific qPCR assay for cancer biomarker detection.

**Abstract:** Analysis of circulating cell-free DNA (cfDNA) methylation abnormalities has emerged as a promising strategy for detecting various diseases, including cancer. While single-biomarker approaches may lack sensitivity, comprehensive next-generation sequencing (NGS) can be cumbersome and expensive. Multiplex methylation qPCR assays offer a practical intermediate solution by providing accurate, accessible, and affordable methylation biomarker detection. In this study, we demonstrated a process for developing a multiplex methylation-specific qPCR assay using colorectal cancer (CRC) methylation biomarkers as a case study. Starting with a set of CRC methylation biomarkers, we developed the Multiplex Methylation-specific qPCR (MMqPCR) algorithm to scan differentially methylated regions (DMRs) for methylation-specific PCR primer and probe design. We then established a systematic process to eliminate assay cross-reactions, reduce background noise, and assess multiplex compatibility. Using a low concentration of hypermethylated DNA (0.1%) in a digital analysis, we demonstrated a 95.8% detection rate with the multiplex strategy, significantly outperforming the singleplex approach (47.9%). The multiplex qPCR development principles and automated primer design algorithm presented here provide valuable tools for developing disease screening, detection or monitoring methods based on methylation analysis.

## Introduction

One of the primary mechanisms of cancer involves the inactivation of tumor-suppressor genes through promoter hypermethylation, leading to gene silencing across various cancer types [1, 2]. Numerous methylation changes, particularly promoter hypermethylation, have been identified as promising biomarkers for disease detection, especially cancer [3, 4]. Combined with non-invasive or minimally invasive liquid biopsy strategies, omics analysis, including methylation analysis, has shown great potential for early cancer detection and monitoring [3-5].

Among current clinical applications, cfDNA methylation analysis for colorectal cancer (CRC) screening has demonstrated the most success. Leveraging recent technological advancements, this method has been commercialized [2-4, 6]. While targeted methylation NGS-based strategies, such as Shield^™^, (Guardant Health), and Galleri®(Grail), have shown significant detection in large clinical trials, their high cost and complex workflows limit their accessibility [7]. Alternatively, simple and affordable single methylation biomarker qPCR assays, like Septin-9 (Epigenomics AG) and SDC2 (Creative Biosciences, Genomictree), have demonstrated limited sensitivity and specificity [3, 8].

To address these limitations, a targeted approach using qPCR to detect a small panel of high-quality biomarkers, as exemplified by Cologuard®, offers a balance between sensitivity and accessibility. By leveraging the wide availability of qPCR systems, this approach provides an accurate, affordable, and accessible option for CRC screening [3, 4, 8]. As demonstrated by NGS methylation analysis, interrogating a cluster of CpG islands within a given differentially methylated region (DMR) significantly improves detection accuracy and sensitivity [9].

While free guidelines and licensed software for developing singleplex methylation-specific PCR assays are available online, few tools support the design of assays targeting multiple CpG sites within a single primer/probe, let alone multiplex qPCR assays. Most current programs for methylation-specific PCR primer design are limited to single CpG site or singleplex format [4]. In this study, we used CRC as a model to demonstrate a new system for automated primer/probe design that integrates multiple CpG sites (more than 3) per DMR and optimizes multiplex methylation-specific qPCR assays in a single reaction.

There are numerous PCR strategies available to analyze methylation-related biomarkers such as methylation-specific restriction endonucleases (MSRE) analysis, methylation-specific high-resolution DNA melting (MS-HRM), and quantitative methylation-specific polymerase chain reaction (qMSP). Each of these methods have advantages and limitations [4, 10], and neither have the same capacity for a multiplexed analysis within one reaction using a common qPCR instrument. The 5’ nuclease qPCR assay can be designed in multiplexing format to interrogate multiple CpGs and maximize the sensitivity and specificity[10].

The system presented here has the potential to facilitate the development of methylation-based multiplex qPCR assays for detecting other cancer types or diseases associated with altered methylation status [2, 11-13].

## Materials and Methods

### DNA samples

Two types of genomic DNA were used for initial primer screening, assay optimization, and final digital testing: 1) Hypermethylated DNA: Human HCT116 DKO DNA (Zymo #D50142-2) and 2) Hypomethylated DNA: Coriell gDNA NA17360 (hypomethylation status confirmed by a previous hybridization-based NGS analysis using a comprehensive methylation oncology panel (Agilent #9419A and #5280-0058)).

Both DNA samples were quantified using a Thermo Fisher Nanodrop 2000 Instrument.

### Bisulfite conversion

Bisulfite conversion was performed using the Zymo EZ DNA Methylation-Lightning^™^ Kit (Zymo #D5030) according to the manufacturer’s instructions. 100-500 ng of DNA was used as input for each conversion reaction.

### qPCR assays

#### Two types of qPCR assays were used in the development process

Initial primer screening and assessment of assay performance and multiplex compatibility were performed using DNA chelating-dye-based qPCR assays. 10 ng of bisulfite-converted DNA input were used in 10 µL qPCR reactions with 2x EvaGreen qPCR MasterMix with ROX dye (Biotium, Cat#31042). Each reaction contained 200 nM of both forward and reverse primers. Primers were synthesized by Integrated DNA Technologies, Inc (IDT).

#### 5’ nuclease qPCR assay

10 ng of bisulfite-converted DNA input were used in a 10 µL reaction with 2x Biotium Universal Probe qPCR Master Mix with ROX dye (Biotium #31044). Each reaction contained 500 nM of each primer and 200 nM of the probe (probe concentration was reduced to 100 nM for multiplex qPCR assays). In the final multiplex assay, probes for every biomarker were FAM dye-labeled. A GAPDH assay with a SUN dye-labeled probe was added as an endogenous control. All probe oligos were synthesized by IDT.

A two-step qPCR protocol was performed on an Applied Biosystems StepOne^™^ qPCR instrument. The protocol consisted of a 2-minute enzyme activation step at 95°C, followed by 40 cycles of denaturation at 95°C for 5 seconds and annealing/extension at 60°C for 30 seconds. Amplification curves were analyzed using an auto-baseline method, and Cq values were calculated using StepOne software by manually setting a threshold line at 0.2 within the exponential phase of amplification across all samples.

A no-template control (NTC) was set up using nuclease-free water.

### Digital analysis of PCR assay

To compare the detection power of multiplex and singleplex PCR assays for a single DNA copy, a digital PCR analysis was performed. A total reaction volume of 480 µL containing 2 ng of bisulfite-converted Zymo’s hypermethylated DNA was distributed across 48 wells of a 5’ nuclease qPCR assay. The number of positive wells was used to calculate the detection rate.

## Results and discussion

### Selection of Target Genes/CpG Islands and sequence conversion

A set of 12 genes—SEPT9, SDC2, SFRP1, SFRP2, PRIMA1, BCAT1_1, BCAT1_2, IKZF1, IRF4, ALX4, OSMR, and VIM—previously identified as CRC epigenetic biomarkers [3-5, 10] were selected. The promoter region sequences of these genes, including all possible CpG islands, were downloaded from the UCSC Genome Browser [14]. To efficiently design primers and probes for multiplex methylation qPCR assays, we developed an automated primer and probe design program: Multiplex Methylation-specific qPCR (MMqPCR).

MMqPCR first performs bisulfite conversion of the input sequence *in silico* to convert cytosines in non-CpG sites into uracils, while maintaining the cytosines in CpG sites. So the primers selected by the MMqPCR program will only amplify unconverted cytosine. This design will ensure that the PCR tests will only amplify the methylated CpG sequences as all the CRC epigenetic biomarkers chosen in this assay are hypermethylated in CRC patients.

Next, the converted promoter region sequence was scanned for possible assay design windows.

### Methylation-specific PCR primer and probe design

After converting the CpG island, the MMqPCR program scans the sequence to identify a design window of less than 150 bp (preferably 60-100 bp) to set the 3’ end of the PCR primers. The primer’s length can be further fine-tuned, while ensuring the following criteria are met: 1) overall GC content between 35% and 65%, 2) at least two CpGs per primer and one CpG in the amplicon insert region, and 3) a 3’ cytosine of a CpG at the end of each primer (Figure 1).

**Figure 1:**
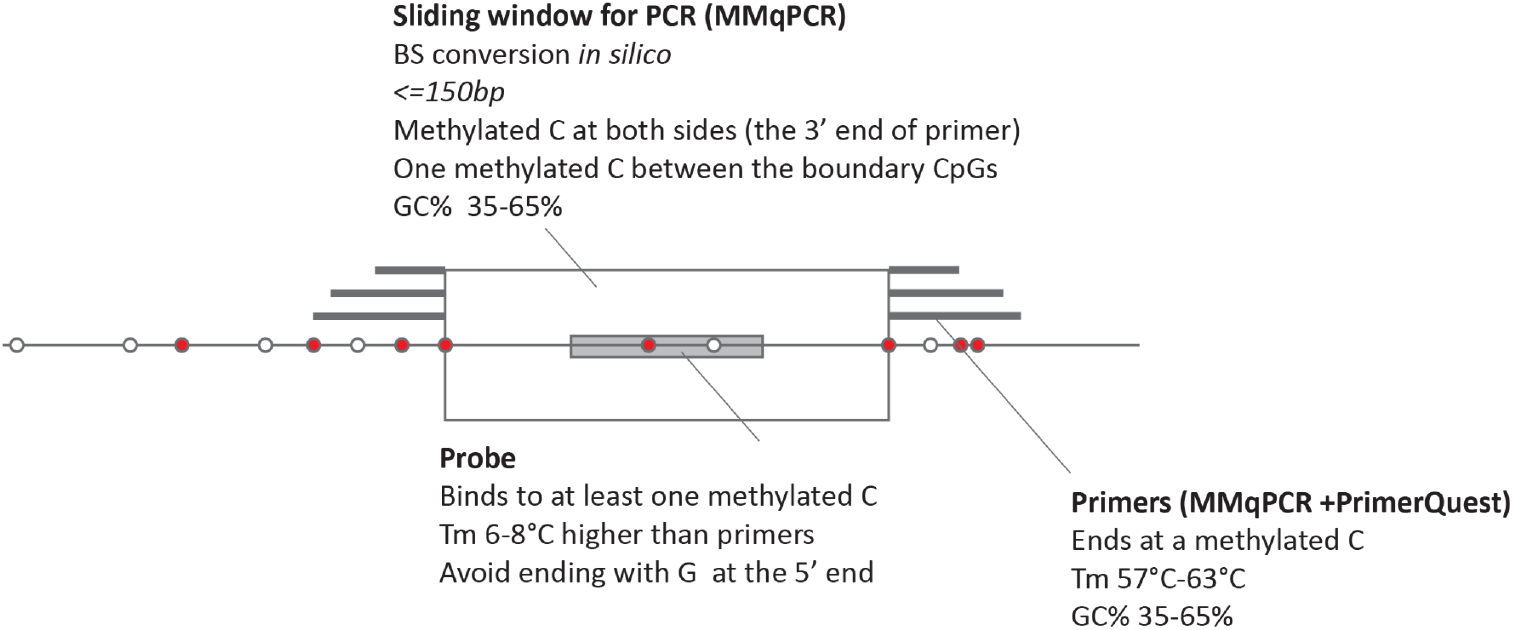
In silico Bisulfite Conversion and Primer/Probe Design. A schematic illustration of in silico bisulfite conversion, PCR amplicon selection window, and primer/probe design is shown. Unmethylated cytosines (non-CpG) are represented by white circles, while methylated cytosines (CpG) are shown as red circles. The PCR amplicon selection window is depicted as a rectangle, primers as gray bars flanking the window, and the probe as a dark gray box.

After using the MMqPCR program to select a design window, we uploaded the output sequences to IDT’s PrimerQuest tool, a publically available online qPCR primer design tool, to design a 5’ nuclease-based qPCR assay. Assays were designed based on the following criterias:

1. Primers should have a melting temperature (Tm) of approximately 57 °C - 63 °C (preferably 60°C).
2. GC% for the primers should be between 35% and 65%.
3. Each primer ends on a C of a CpG at the 3’ end.
4. Probes must be between forward and reverse primers.
5. Probes must not overlap with forward primer sequences.
6. Probes must have a melting temperature 6-8°C higher than that of the primers.
7. There must be at least 1 CpG in the probe, preferably in the center.
8. Probes should not end with a G at the 5’ end to prevent dye quenching.

Primer sequences identified by MMqPCR were re-entered into the program for probe sequence selection and evaluation. Assays with the following issues were eliminated:

1. Four guanines (GGGG) in a row
2. Two sets of GGG in a row within 8 or less nucleotides
3. Primer has high mapping hits in BS converted genome (>100)
4. Difference in temperature of probe and primer is not within 6-8 °C

One candidate assay for each biomarker was selected for next step analysis.

Code for MMqPCR program is deposited at https://github.com/hello2632/MMqPCR/blob/541142f1ac97eda82bb71177576da009a04f9609/test_version.ipynb

### Singleplex EvaGreen qPCR assay testing

Assays were formulated using only primer oligonucleotides (Table 1) and tested with a DNA chelating-dye-based qPCR system. Assays meeting the following criteria were selected as candidate assays for multiplexing:

**Table 1:**
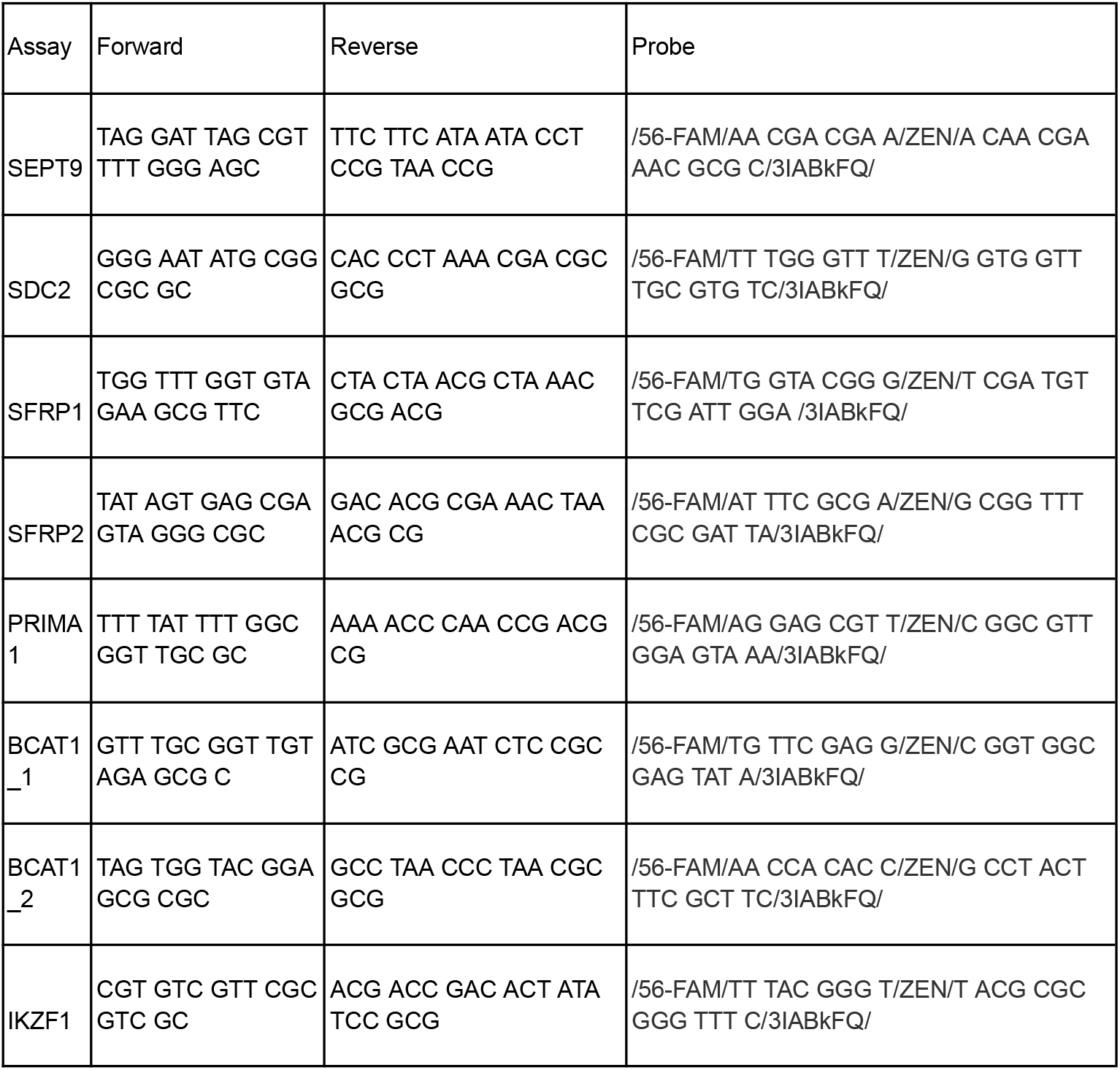
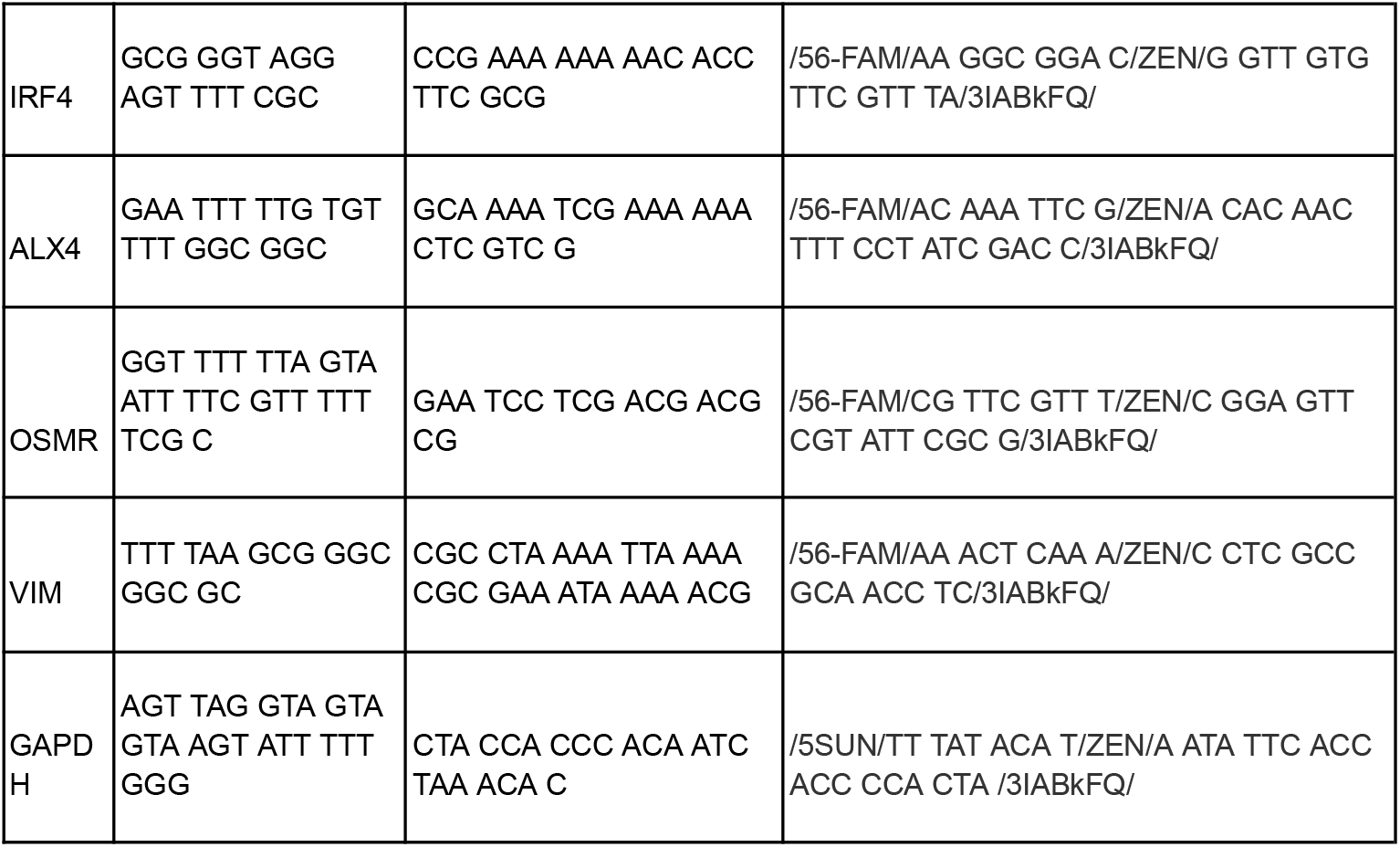
Primer and Probe Sequences for 12 Methylation Assays and the GAPDH Control Assay. All 12 methylation assays use FAM-labeled probes with ZEN/Iowa Black™ FQ quenchers. The GAPDH control assay uses a SUN-labeled probe with a ZEN/Iowa Black™ FQ quencher.

1. Efficient amplification with bisulfite-converted Zymo’s HCT116 hypermethylated DNA (Cq < 32 at 10 ng gDNA input).
2. Little to no amplification with bisulfite-converted hypomethylated gDNA Coriell NA17360 to prevent background amplification in normal samples (Cq > 38 at 10 ng gDNA input).
3. No amplification with no template control.

10 out of the 12 assays successfully met the above criteria.

### Duplex EvaGreen qPCR assay testing

Following the identification of each potential assay, we formulated duplex assays of every combination of the selected singleplex assays using DNA chelating-dye qPCR assays. A single DMR assay can be considered for the final multiplex assay based on the following criteria for each duplex assay:

1. Efficient amplification with bisulfite-converted Zymo’s HCT116 hypermethylated DNA (Cq < 32 at 10 ng gDNA input)
2. Little to no amplification with bisulfite-converted hypomethylated gDNA Coriell NA17360 (Cq > 38 at 10 ng gDNA input)
3. No amplification with no template control.
4. Compatible with endogenous control GAPDH assay - no amplification when formulated with GAPDH assay in duplex format.

Primer-dimer interactions were experimentally characterized. The data on failed duplex combinations was used to calibrate the MMqPCR program, enhancing its capacity to identify potential primer-dimer formation within and between individual assays during multiplex assay design.

### Single plex 5’ nuclease qPCR assay testing 5’ nuclease qPCR assay

Probes were added to the selected primers to form singleplex 5’ nuclease qPCR assays. Assays meeting the following criteria were selected as candidates for multiplex assay development:

1. Efficient amplification with the bisulfite-converted Zymo’s HCT116 hypermethylated DNA (Cq < 32 at 10 ng gDNA input)
2. No amplification with the bisulfite-converted hypomethylated gDNA Coriell NA17360 to prevent background amplification in normal samples (Not detected at 10 ng gDNA input)
3. No amplification with no template control.

### Multiplex 5’nuclease qPCR assay analysis

Based on different combinations of compatible duplex assays, two sets of four-plex 5’-nuclease qPCR assays were formulated:

Assay A: A 4-plex FAM probe assay including SEPT9, IRF4, VIM, and BCAT1_1 (each with a FAM probe), and a control GAPDH assay with a SUN probe.

Assay B: A 4-plex FAM probe assay including SEPT9, IRF4, VIM, and ALX4 (each with a FAM probe), and a control GAPDH assay with a SUN probe.

Both sets of multiplex assays were tested with three DNA templates: bisulfite-converted hypermethylated DNA, bisulfite-converted hypomethylated Coriell gDNA NA17360, and a no-template control. Based on Cq values, Assay B demonstrated slightly higher sensitivity than Assay A for hypermethylated DNA (Table 2). Therefore, Assay B was selected as the final assay.

**Table 2:**
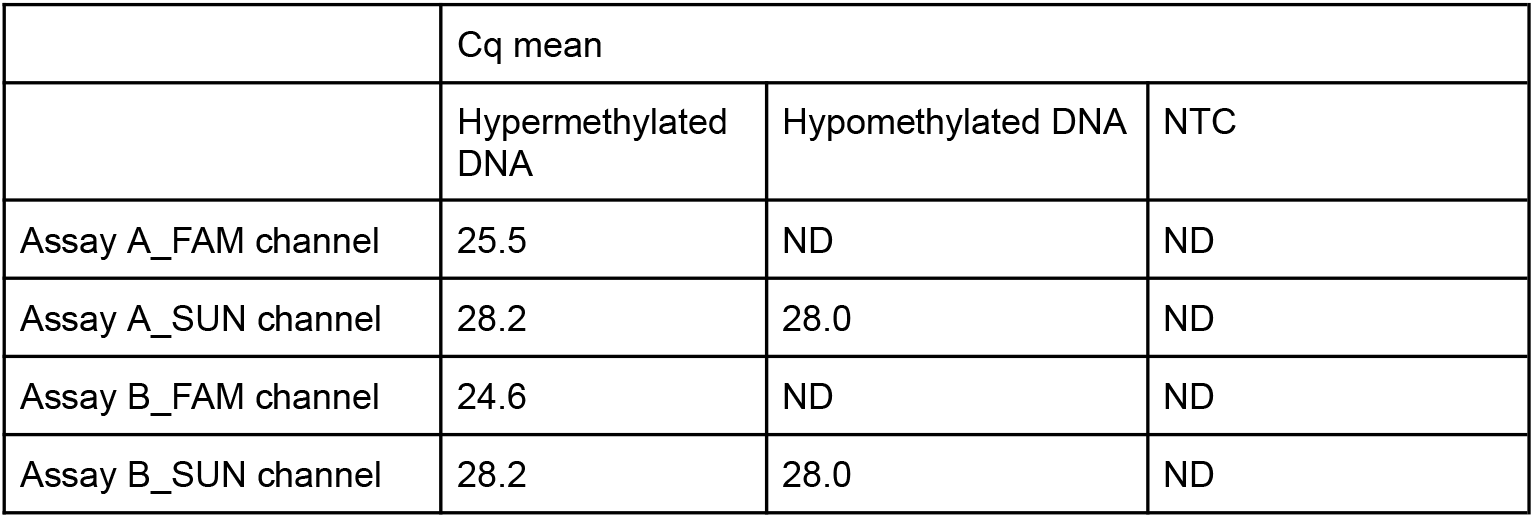
Performance of Two Multiplex Assay Sets. Performance of the assay on positive (hypermethylated DNA), negative (hypomethylated DNA), and no-template control (NTC) samples. The FAM channel reports the accumulated signal of methylation for the four biomarkers in each assay. The SUN channel (VIC channel on the StepOne qPCR instrument) reports the DNA input control (GAPDH). Both assay sets successfully detected positive controls and showed no detection (ND) for negative controls and NTC. The DNA input GAPDH control assay showed comparable detection for both positive and negative controls.

|  | Cq mean |  |  |
| --- | --- | --- | --- |
|  | Hypermethylated DNA | Hypomethylated DNA | NTC |
| Assay A_FAM channel | 25.5 | ND | ND |
| Assay A_SUN channel | 28.2 | 28.0 | ND |
| Assay B_FAM channel | 24.6 | ND | ND |
| Assay B_SUN channel | 28.2 | 28.0 | ND |

### Analytical analysis of 4-plex assay

#### 4-plex vs. 1-plex SEPT9 assay comparison

The developed 4-plex assay (SEPT9, IRF4, VIM, ALX4 with GAPDH) was compared to a singleplex SEPT9 assay (with GAPDH). SEPT9 was chosen as the singleplex benchmark because it is the first FDA-approved methylation biomarker and the most widely used single biomarker for CRC pre-screening (Epi proColon). Both assays were tested against three templates: bisulfite-converted hypermethylated DNA, hypomethylated DNA, and a no template control (Figure 2). The 4-plex assay demonstrated approximately 2 cycles lower Cq values compared to the SEPT9 singleplex assay, indicating significantly higher sensitivity.

**Figure 2:**
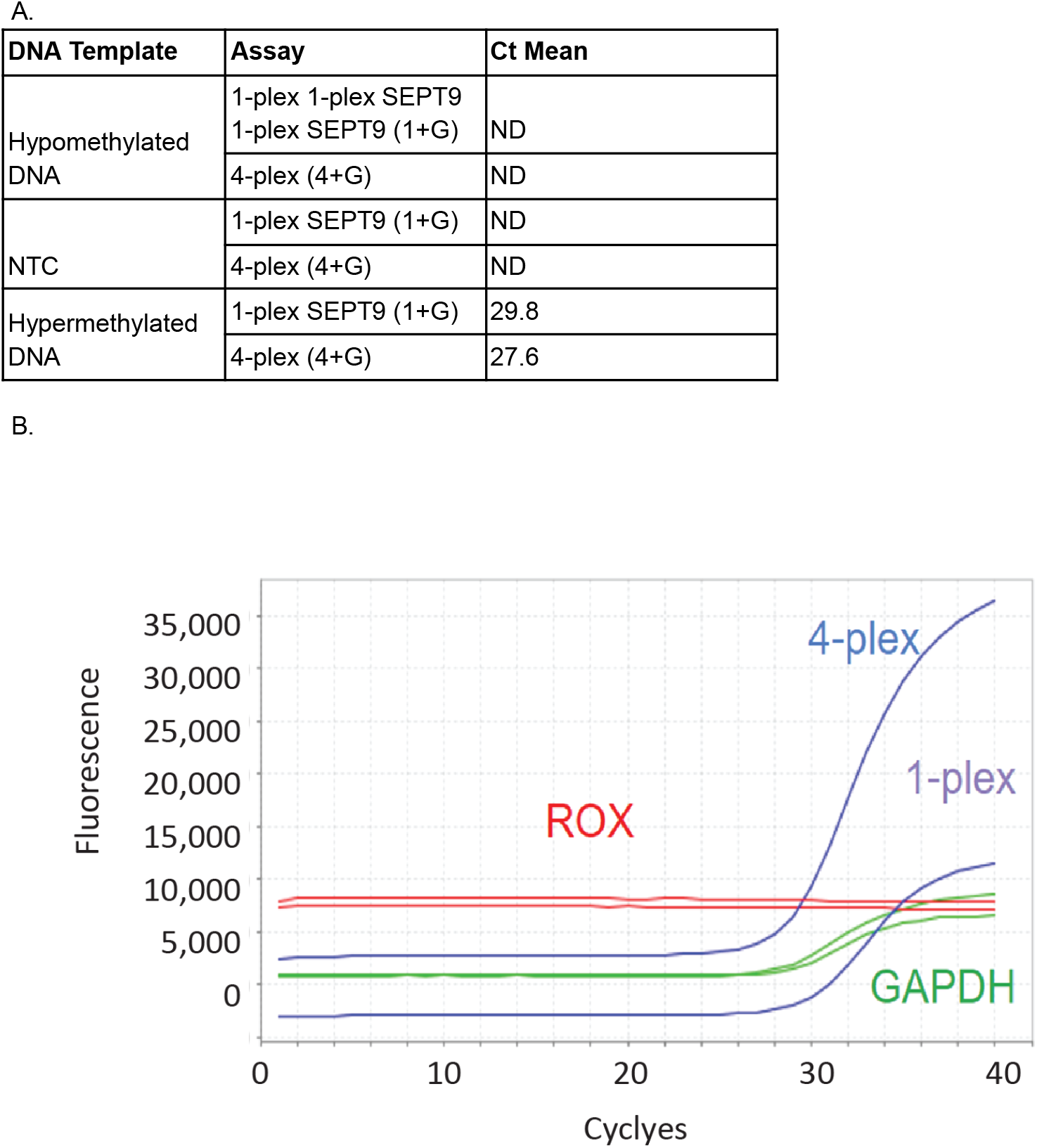
Comparison of 4-plex and Singleplex Assay Performance. **A.** The 4-plex assay (4+G) shows a higher accumulated methylation signal (indicated by a lower mean Cq value) compared to the singleplex assay (1+G) for the positive control (hypermethylated DNA). **B**. qPCR readout of the 4-plex (blue) and 1-plex (purple) assays on the positive control (hypermethylated DNA). ROX (red) is used as a reference dye, and GAPDH (green) is used to report DNA input quantity and normalize methylation detection.

### Assay sensitivity

To analyze the ability of the 4-plex assay to detect low levels of methylation, we spiked Zymo’s HCT116 hypermethylated DNA into Coriell NA17360 hypomethylated DNA at various percentages (10%, 1%, 0.1%, 0.03%, and 0%). The DNA mixtures were bisulfite-converted, and 10 ng of DNA was used per reaction. The 4-plex assay demonstrated a higher sensitivity, detecting hypermethylated DNA down to 0.03% compared to the singleplex assay, which could only detect down to 0.1% (Figure 3).

**Figure 3:**
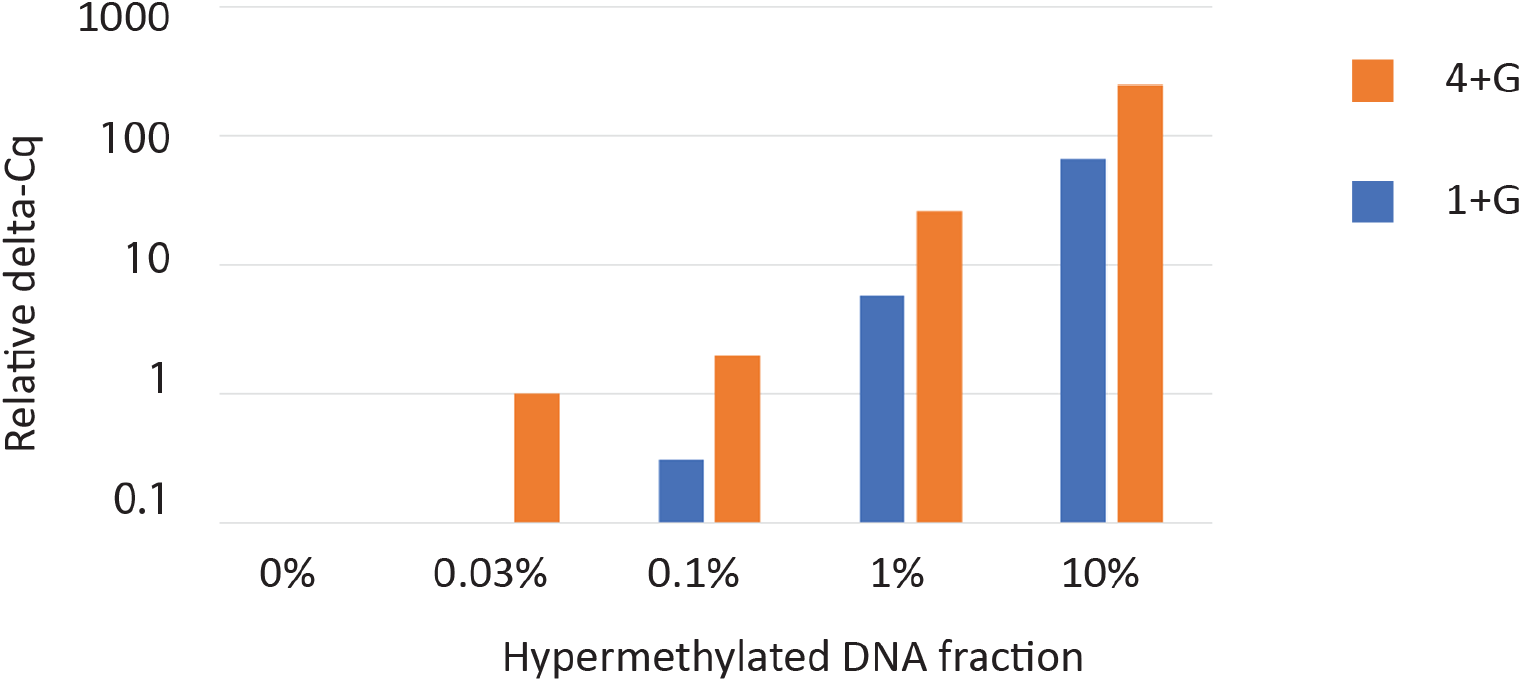
Relative Quantification of 4-plex and Singleplex Assays. This figure shows the relative quantification based on delta-Cq values. The Cq value of the 4+G assay on the 0.03% spike-in sample was set as the baseline for quantification (Quant = 1), and other assay readings were compared to this baseline to derive relative quantification. A cutoff of 0.1 times the baseline was set, and the Cq values of the 0% sample and the 1+G assay on the 0.03% spike-in sample were below this cutoff and not shown. The 4-plex assay (4+G, orange) demonstrates a higher detection rate (log-based) compared to the singleplex assay (1+G, blue) across different fractions of hypermethylated DNA in an unmethylated DNA background.

### Digital analysis of assay sensitivity

To further demonstrate the superior sensitivity of the 4-plex assay over the singleplex assay, we conducted a 48-well digital analysis experiment, using Poisson distribution to analyze each assay’s ability to amplify a single DNA copy.

To simulate the low tumor content in low quantity of possible blood cfDNA from clinical samples, hypermethylated DNA was spiked into hypomethylated DNA at a 0.1% concentration in a total of 2ng input (666 copies of genome). The DNA mixture was then bisulfite-converted. To test the detection limit of the multiplex assay, we used 2 ng of the 0.1% DNA mixture as the input per well for 48-well test, resulting in a theoretical average 0.67 copies of the methylated genome per well.

Based on Poisson distribution calculation, the singleplex assay was predicted to yield approximately 23.4 positive wells (λ = 0.67, P(X≥1) = 48.8%), assuming 100% efficiency in detecting target molecules. Similarly, the 4-plex assay was predicted to yield approximately 44.7 positive wells (λ = 2.68, P(X≥1) = 93.1%) in the 48-well assay.

The SEPT9 singleplex (1+G) assay amplified 23/48 wells (47.9% sensitivity), closely matching the theoretical 48.8%. The 4-plex assay amplified 46/48 wells (95.8% sensitivity), also aligning with the theoretical 93.1%, demonstrating the robustness of the 4-plex assay system (Figure 4). Compared to the singleplex SEPT9 assay, a reference for current clinical qPCR tests, the multiplex assay offers significantly improved theoretical and observed sensitivity.

**Figure 4:**
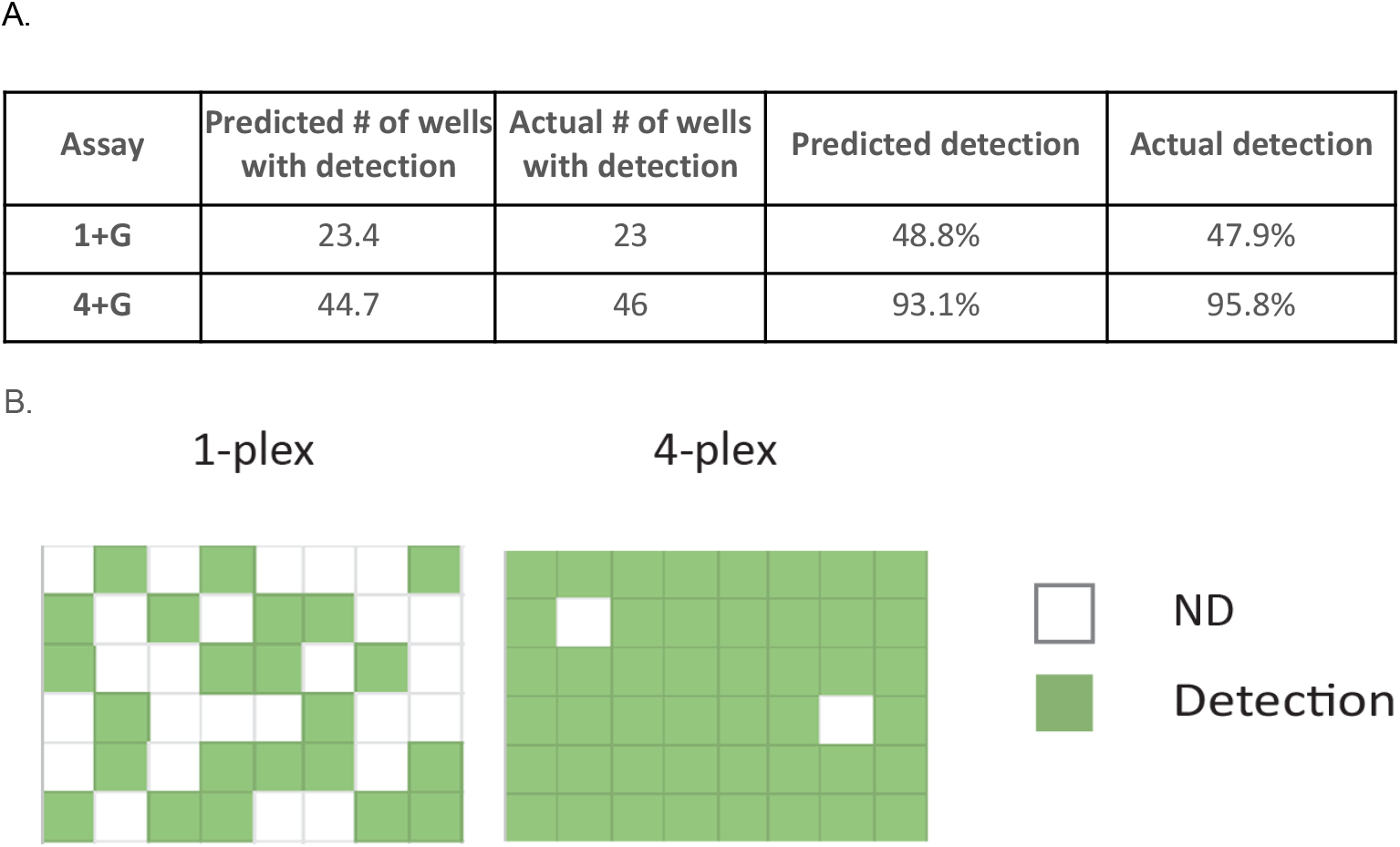
Digital PCR Analysis of 1-plex and 4-plex Assays. **A.** Proportion of positive wells (with DNA detection) out of 48 wells in the digital PCR experiment. **B**. Distribution of DNA detection across the 48-well digital PCR plate.

## Conclusion

To develop an accessible and robust method for multiplex methylation qPCR assays, we created the MMqPCR program to automate primer and probe design. Using hypermethylated colorectal cancer (CRC) biomarkers as a model, we formulated a multiplex qPCR assay that demonstrated the increased sensitivity of a multiplex assay compared to a singleplex assay.

For future improvements, we plan to expand the MMqPCR program to analyze inter-assay primer interactions. Additionally, we aim to develop higher-plex assays capable of detecting even lower concentrations of methylated genomes while maintaining the sensitivity of our four-plex assay. Ultimately, our goal is to refine the multiplex assay design algorithm to enable higher plexity, resulting in more sensitive, accessible, and efficient assays for disease detection.

